# demuxSNP: supervised demultiplexing scRNAseq using cell hashing and SNPs

**DOI:** 10.1101/2024.04.22.590526

**Authors:** Michael P. Lynch, Yufei Wang, Laurent Gatto, Aedin C. Culhane

**Affiliations:** Limerick Digital Cancer Research Centre, Health Research Institute, School of Medicine, University of Limerick, Ireland; Department of Cancer Immunology and Virology, Dana-Farber Cancer Institute, Boston, MA, 02215, USA; Harvard Medical School, Boston, MA, 02115, USA; Computational Biology and Bioinformatics Unit (CBIO), de Duve Institute, UCLouvain, Belgium

**Keywords:** Single-cell, demultiplexing, cell hashing, SNPs

## Abstract

**Background:** Multiplexing single-cell RNA sequencing experiments reduces sequencing cost and facilitates larger scale studies. However, factors such as cell hashing quality and class size imbalance impact demultiplexing algorithm performance, reducing cost effectiveness

**Findings:** We propose a supervised algorithm, demuxSNP, leveraging both cell hashing and genetic variation between individuals (SNPs). The supervised algorithm addresses fundamental limitations in demultiplexing with only one data modality. The genetic variants (SNPs) of the subset of cells assigned with high confidence using a probabilistic hashing algorithm are used to train a KNN classifier that predicts the demultiplexing classes of unassigned or uncertain cells. We benchmark demuxSNP against hashing (HTODemux, cellhashR, GMM-demux, demuxmix) and genotype-free SNP (souporcell) methods on simulated and real data from renal cell cancer. Our results demonstrate that demuxSNP outperformed standalone hashing methods on low quality hashing data, improving overall classification accuracy and allowing more high RNA quality cells to be recovered. Through varying simulated doublet rates, we show genotype-free SNP methods are unable to identify biological samples with low cell counts at high doublet rates. When compared to unsupervised SNP demultiplexing methods, demuxSNP’s supervised approach was more robust to doublet rate in experiments with class size imbalance.

**Conclusions:** demuxSNP is a performant demultiplexing approach that uses hashing and SNP data to demultiplex datasets with low hashing quality where biological samples are genetically distinct. Unassigned cells (negatives) with high RNA quality can be recovered, making more cells available for analysis, especially when applied to data with low hashing quality or suspected misassigned cells. Pipelines for simulated data and processed benchmarking data for 5-50% doublets are publicly available. demuxSNP is available as an R/Bioconductor package (https://doi.org/doi:10.18129/B9.bioc.demuxSNP).

## Introduction

Single-cell RNA sequencing (scRNAseq) enables insight into cellular heterogeneity, cell sub-types and cell-cell communication not previously possible with bulk methods due to gene expression averaging [1]. Cost remains a barrier for large scale research and clinical studies at a single-cell resolution [2] despite reductions in cost of sequencing technologies. Multiplexing in scRNAseq refers to the sequencing of cells from multiple different biological samples on the same sequencing lane, rather than on individual lanes. This reduces sequencing costs and technical batch effects [3]. The cells must then be demultiplexed, or assigned back to their biological sample of origin prior to downstream analysis. In droplet-based technologies, higher cell loading rate results in a higher doublet rate, thus limiting the lane capacity. In multiplexed experiments, doublets made up of cells from different samples are more easily identified and removed, allowing higher cell loading rate onto the sequencing lane. Demultiplexing strategies can be broadly divided into two approaches, experimental cell tagging (cell hashing) and bioinformatics analysis of genetic variation using single nucleotide polymorphisms (SNPs). Cell hashing is popular due to its applicability to a wide variety of experimental designs and commercially available kits. SNPs based methods are limited to genetically distinct samples but have lower library preparation costs.

Cell hashing is a combined experimental and computational approach where cells from each biological sample are labelled with a distinct sequenceable tag [4,5] prior to being pooled and sequenced. Computational algorithms, such as those reviewed by Howitt et al. [6], then operate on the resulting counts matrix to determine which cells are associated with which biological sample of origin. However, technical artefacts such as non-specific binding, doublets and varying cell quality complicate this procedure. In our experience, cell line or blood single cell studies largely have higher hashing quality, however studies of solid tumour tissue where processing of cells involves complex dissociation protocols can reduce hashing quality. Cells with low hashing quality may be assigned to the incorrect group. Additionally, cells deemed to have no hashing signal in any group are referred to as hashing negatives, or negatives for short. Small numbers of hashing negatives are permissible, however, large numbers of negatives result in wasted data and so are undesirable. Due to their dependence on hashing quality, performance of standalone hashing based demultiplexing methods can vary significantly between datasets [7].

SNP based demultiplexing methods exploit natural genetic variation between genetically distinct biological samples. Genotype based methods such as Demuxlet [8] and scSNPdemux [9] require a priori knowledge of the genotype of each biological sample, incurring additional experimental cost and limiting their utility. Genotype-free methods [10–12] are more commonly used but face their own unique limitations. While they can group cells, they cannot link cells back to a biological sample without additional genotype or hashing data. Calling SNPs in scRNAseq is challenging as the data is sparse with reads concentrated in specific regions and gene expression can be highly variable within a dataset [13]. Performance is dependent on the sequencing depth and reduces in datasets with high levels of ambient RNA [14] or in cells such as cancer cells, where somatic mutation can obscure the germline signal. Despite the considerable number of methods developed, a universally robust tool has yet to be developed.

Hashing negative cells which cannot be assigned are removed prior to downstream analysis steps resulting in wasted data. Additionally, researchers also exclude cells if there is disagreement between demultiplexing algorithms or low probability assignment. This results in further wasted data and reduces the effectiveness of multiplexing as a cost-saving measure. Alternatively, retaining uncertain cells which may be wrongly assigned reduces the statistical power of differential gene expression analysis and confounds biological interpretations in downstream analysis steps.

We propose a supervised multi-modal method, demuxSNP, that leverages both SNP and hashing modalities. We benchmark against existing standalone hashing (HTODemux [4], BFF_raw and BFF_cluster [15], GMM-Demux [16], demuxmix [17]) and genotype-free SNPs based (souporcell [12]) methods, adapting existing SNP simulation pipelines [14] paired with hashing data, to better understand performance across a range of scenarios against reliable ground truth. We further motivate the utility of demuxSNP over popular existing methods with application to a case study dataset. demuxSNP is available as an R/Bioconductor package.

## Results

### 1. Overview of demuxSNP

A key challenge in hashing based demultiplexing is variability in hashing quality due to technical issues such as non-specific binding. In general, a proportion of cells from each group may be confidently called, the number of which will depend on the hashing quality for a specific experiment (Figure 1A). These high confidence cells may be identified using consensus methods such as cellhashR [15], probabilistic methods with high acceptance threshold [16,17], or use of non-conservative count threshold to describe the positive peak, however, retaining only these high confidence cells will result in loss of valuable data.

**Figure 1.**
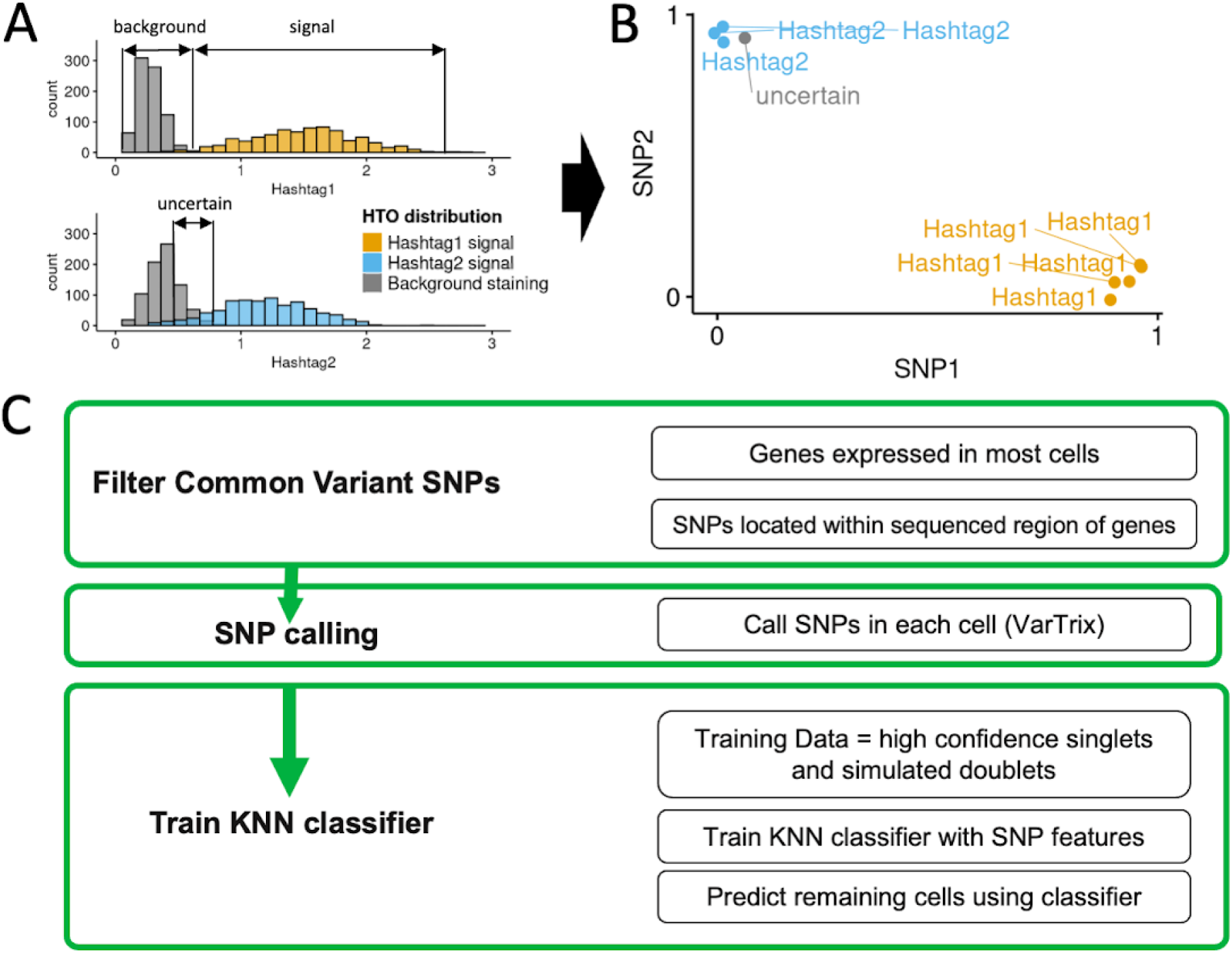
Overview of the demuxSNP workflow. (A) high quality hashtag counts can be separated into a bimodal distribution with distinct signal and background peaks. Low quality hashtag counts have a poorly separated bimodal distribution and have high numbers of misassigned, uncertain or hashing negative cells. (B) SNP profiles of cells called with high confidence using hashing based methods may be used to predict classes of cells which could not be called with high confidence (uncertain/negative) using hashing methods. (C) demuxSNP workflow involves filtering of SNPs, calling SNPs in each cell then assigning uncertain or hashing negative cells based on SNP data from cells called with high confidence.

For the cells which can not be confidently called using hashing methods, we propose that their correct group may be more easily identified based on their SNP profile (Figure 1B). Training a classifier on the SNP profiles of the high confidence singlet cells and simulated doublets, we predict the class of negative or uncertain cells. We apply demuxmix [17], a highly performant probabilistic demultiplexing algorithm to hashing counts data to determine which cells can be confidently called. We then use demuxSNP to train a classifier to predict the labels of the remaining cells based on their SNP profiles. With high quality hashing data, often a large proportion of cells can be called with high confidence. With low quality hashing data, non-trivial numbers of cells may be assigned as negative or uncertain and their recovery warranted using a method such as demuxSNP. The demuxSNP workflow is outlined in Figure 1C.

### 2. Demultiplexing performance improves when using demuxSNP compared to standalone hashing methods on datasets with poor hashing quality

Benchmark data is simulated from a multiplexed experiment with six hashtags from genetically distinct samples. Aligned reads and hashing counts from singlets assigned with high-confidence by demuxmix [17] are retained. Doublets are simulated from the singlet data by randomly renaming barcodes on aligned reads [14] and summing counts across singlet cells comprising each doublet for SNP calling and hashing data respectively. Features associated with high quality hashing include well separated bimodal peaks, high signal to noise ratio (Figure 2A). Other experimental factors that may improve demultiplexing performance include well balanced group sizes. Hashing quality is reduced by scaling down the signal in each hashtag group. Poor hashing quality is then associated with features such as poor peak separation and low signal to noise ratio, with high class imbalance also impacting demultiplexing performance (Figure 2B).

**Figure 2.**
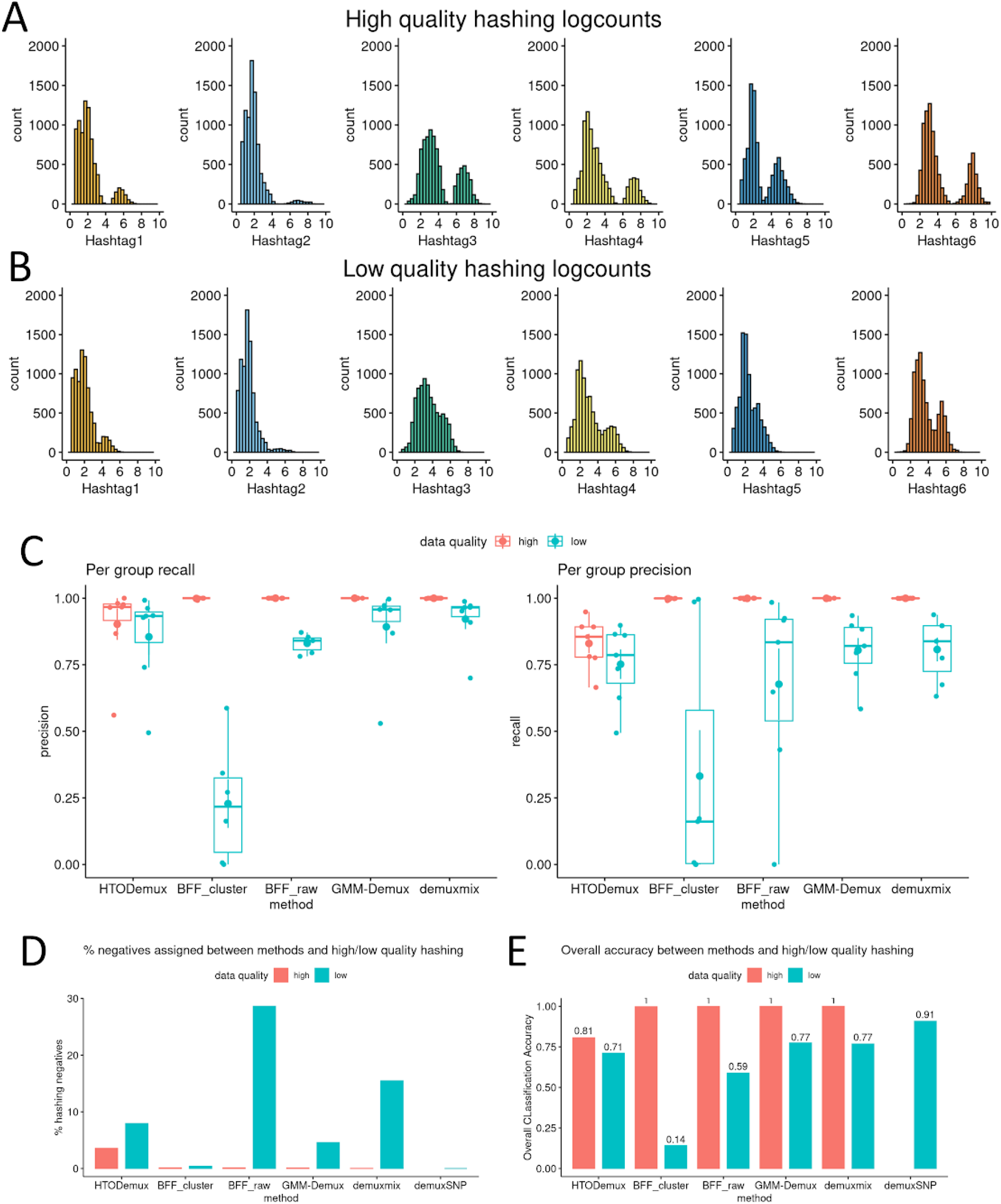
demuxSNP improves cell assignment on datasets with low hashing quality. (A) In benchmarking high quality hashing, the signal and background are distinct. (B) In benchmarking low quality hashing, there is poor separation between signal and background. (C) Hashing algorithm performance decreases with hashing quality. (D) Low quality hashing may result in large numbers of hashing negative cells. (E) demuxSNP shows increased overall classification accuracy on low quality hashing data compared with standalone hashing methods.

Demultiplexing performance is dependent on the quality of hashing. When we compare performance of several popular hashing based algorithms we observe that the performance decreases regardless of method when applied to low quality simulated hashing data with typical doublet rate (20%). On the high quality dataset, HTODemux shows poorest performance compared to the other methods tested for both precision and recall, potentially due to features other than peak separation such as imbalance in class sizes and the misalignment of the signal peaks (Figure 2C). Other methods BFF_raw, BFF_cluster, GMM-Demux and demuxmix each show almost perfect precision and recall on the high quality dataset, an expected result given the clear separation between signal and background. On the low quality dataset, BFF_raw performs poorly, potentially due to the assumption of a bimodal distribution, the extent of which is reduced in this test case.

We explored which methods recovered more cells and thus had fewer cells with no identity (hashing negatives). In terms of the number of assigned hashing negatives (Figure 2D), on the high quality dataset HTODemux assigns the most negatives (∼3.5%) although this falls within an acceptable range. A negligible number of negatives are assigned by BFF_raw, BFF_cluster, GMM-Demux and demuxmix (<0.2%). On the low quality dataset, most negatives are assigned by BFF_raw and demuxmix (28% and 15% respectively). demuxSNP avoids the classification of hashing negatives by leveraging SNP data to assign these cells, reducing wasted data. Fewest cells are assigned as hashing negative by BFF_cluster, however, we note that while high numbers of hashing negatives are undesirable, this reflects only one aspect of algorithm performance and must be taken in context of other classification performance metrics.

We next looked at overall classification accuracy (Figure 2E). Each of the stand-alone hashing algorithms, with the exception of HTODemux, perform almost perfectly on the high quality dataset. We do not compare demuxSNP on this dataset as a large number of cells (∼99%) have already been confidently called by standalone probabilistic hashing algorithms, and thus the use of demuxSNP is not warranted. On the low quality dataset, performance is comparable between HTODemux, BFF_raw, GMM-demux and demuxmix. Despite assigning fewer negatives, BFF_cluster has the lowest accuracy, potentially due to the assumption of a bimodal distribution being challenged here. We show that performance improves when using SNPs to reassign uncertain cells compared to hashing classification alone, with overall classification accuracy of 0.91 for demuxSNP compared to 0.77 and 0.77 for the top performing standalone hashing methods, GMM-demux and demuxmix respectively.

### 3. demuxSNP is more robust to class size imbalance compared to genotype-free SNP method souporcell

We next asked how demuxSNP performs against standalone genotype-free SNP based methods. We first compared overall classification accuracy against souporcell, a popular genotype-free SNP based method across a range of doublet rates from 5-50% [18]. On our simulated datasets, demuxSNP outperforms souporcell at higher doublet rates (>30%) (Figure 3A) but souporcell outperforms demuxSNP in terms of both accuracy and adjusted rand index at low doublet rates (<25%). We investigate assignment patterns in more detail below. Due to the SNP selection feature used by demuxSNP, computational cost is reduced by a factor of 2.5 (Figure 3B).

**Figure 3.**
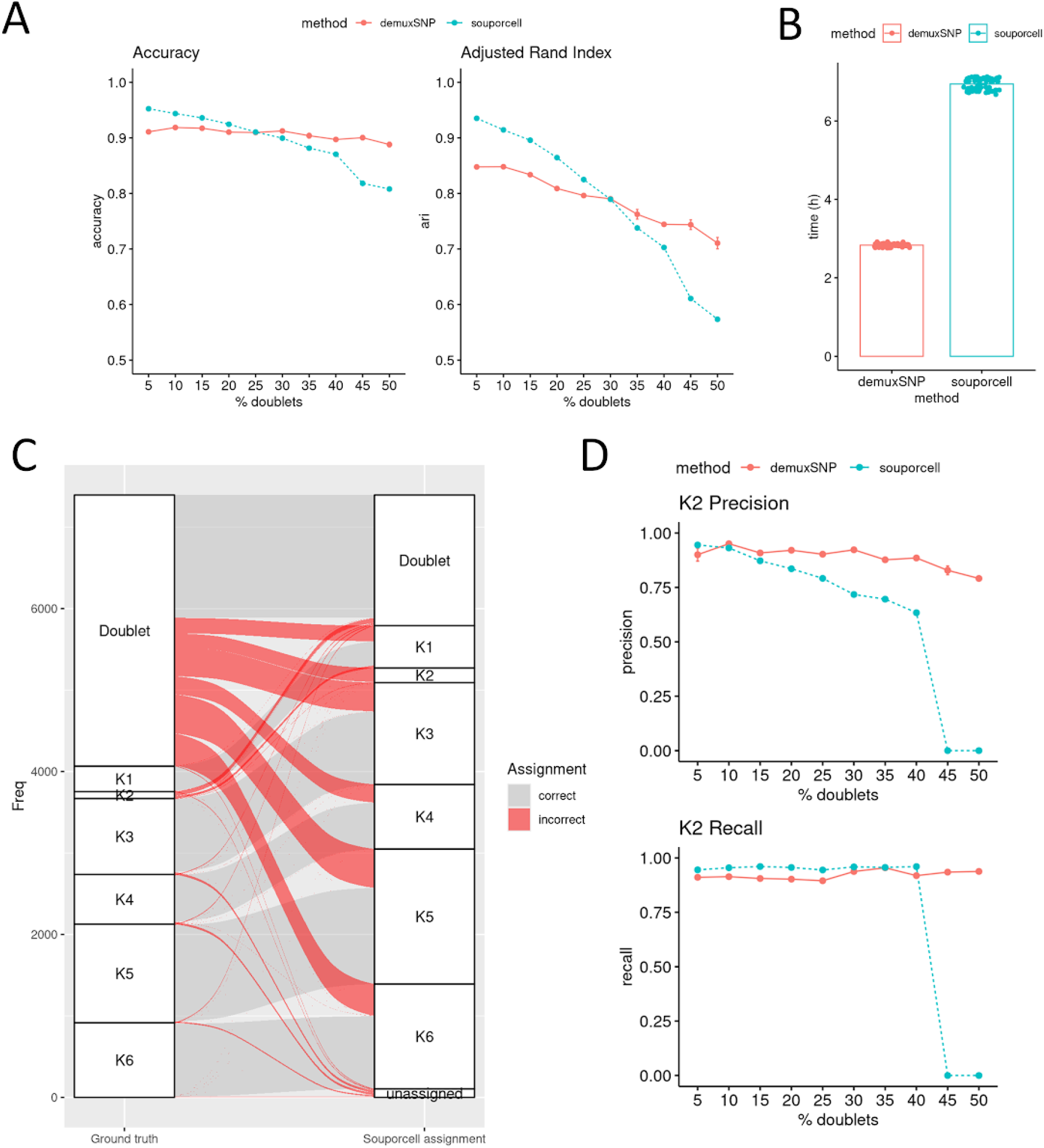
demuxSNP shows increased robustness to class size imbalance compared to genotype-free SNP based method souporcell. (A) demuxSNP (average of 5 runs ± sd) outperforms souporcell (single run, seed fixed) at higher doublet rates. (B) demuxSNP SNP filtering results in reduced computational cost compared to souporcell. (C) Doublets generally overassigned to singlet groups and completely misassigned to K2 (smallest sample) group at 45% doublets. (D) Precision and recall for K2 (smallest group) across a range of doublets is stable for demuxSNP (average of 5 runs ± sd) but reduces to zero past the threshold of 40% doublets for souporcell (single run, seed fixed).

In evaluating the performance of clustering methods on scRNAseq gene expression data, significant attention is given to methods’ ability to detect small clusters [19]. Genotype-free SNPs based methods may face similar challenges, in clustering genetically distinct SNP profiles, where the number of cells per biological sample may vary, intentionally or otherwise. We note that at high doublet rates, souporcell generally assigns large numbers of doublets to singlet groups (Figure 3C). Additionally, the K2 (smallest) group appears to be completely misassigned. The true K2 cells have been assigned as doublets, while the cells assigned as K2 are true doublets. In contrast, demuxSNP correctly identifies the K2 group and assigns fewer doublets (Supplementary Figure 1). The assignment of a large proportion of doublets to a singlet group has the potential to confound downstream analysis if not identified.

To investigate this further, we next asked whether demuxSNP’s supervised approach was more robust in calling biological samples with fewer cells (imbalanced classification) compared to unsupervised methods when doublet rate varies. At lower doublet rates, demuxSNP and souporcell perform comparably. However, at higher doublets rates (>40%), both the precision and recall for souporcell reduce to zero (Figure 3D), whereas demuxSNP’s performance remains stable.

### 4. demuxSNP overcomes demultiplexing challenges in case study dataset

We next demonstrate the utility of demuxSNP on a case study dataset containing cells from six genetically distinct samples from renal cell cancer. We initially identify features in the hashing counts indicating poor quality (Figure 4A) including low signal to noise ratio (Hashtag 3,5), low signal (Hashtag2) and misaligned peaks (Hashtag5). Applying HTODemux (a popular hashing based method), souporcell (a popular genotype-free SNPs method) and demuxSNP, significant differences in results are observed. Notably, HTODemux assigned a large number of hashing negative cells and souporcell shows little agreement with HTODemux and demuxSNP in the Hashtag2 group (Figure 4B).

**Figure 4.**
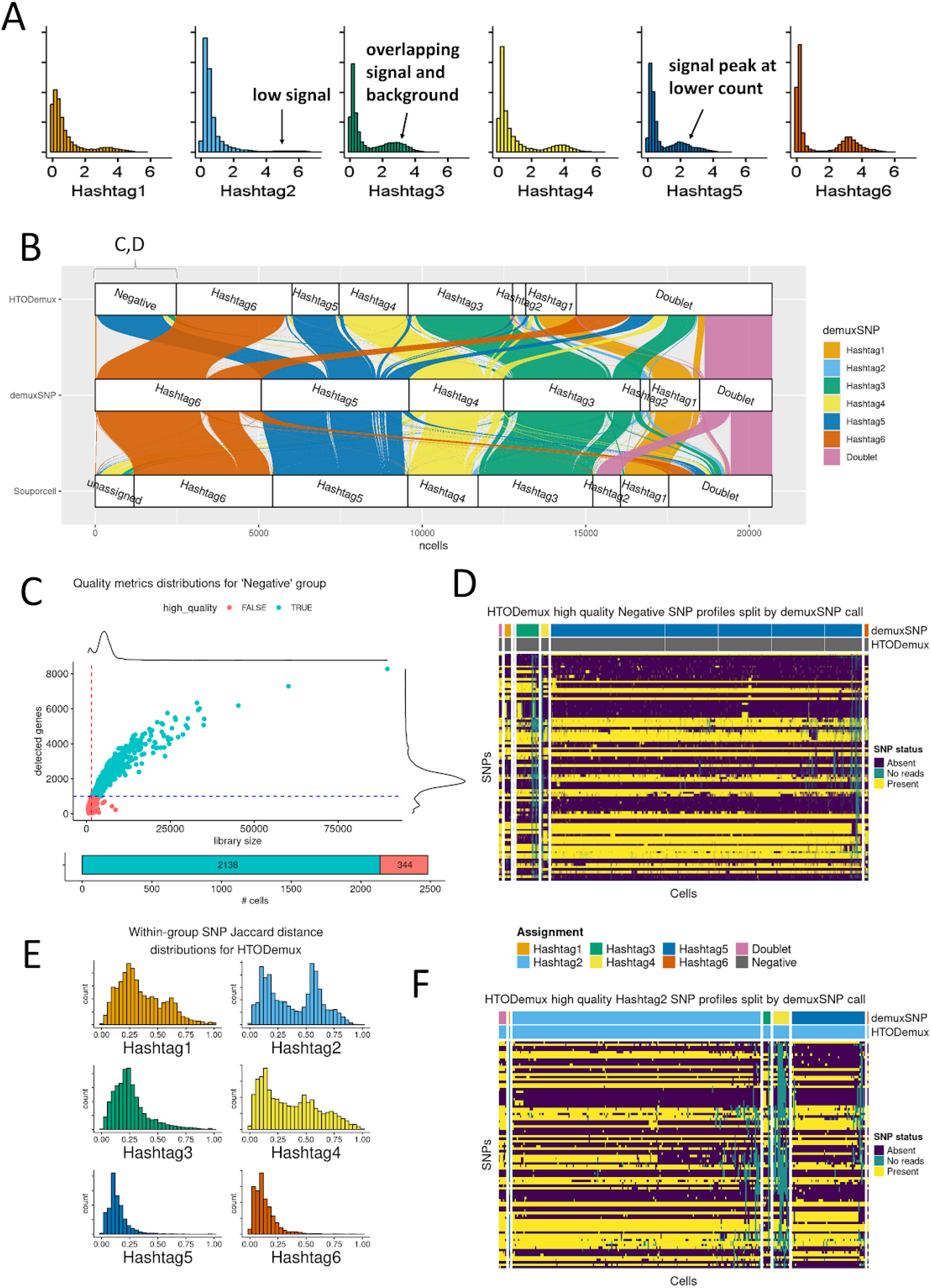
demuxSNP overcomes demultiplexing challenges on real data. (A) Real data contains features of low quality hashing such as low signal, low signal to noise ratio, and unaligned signal peaks. (B) Majority of negatives called by HTODemux assigned as Hashtag5 by demuxSNP and souporcell. Majority of Hashtag2 called by souporcell are called as doublet by demuxSNP and HTODemux. (C) Quality metric distribution for HTODemux negative group. (D) SNP profiles of HTODemux negative group. (E) Bimodal distribution of binary distance matrix for HTODemux singlet group SNP profiles. (F) SNP profiles of HTODemux Hashtag2 group shows multiple SNP profiles.

For souporcell, poor agreement is seen for the Hashtag2 group compared with HTODemux and demuxSNP. The majority of cells assigned as Hashtag2 by HTODemux and demuxSNP have been assigned to Hashtag4 or Doublet group by souporcell. More concerningly, a large number of cells (n=1043) were assigned to the Hashtag2 group which were consistently called as doublets by demuxSNP and HTODemux, leading to significant potential for confounding downstream analysis steps. This is consistent with the behaviour explored in Figure 3C-D whereby souporcell was unable to identify biological samples with small numbers of cells in datasets with high doublet rates. demuxSNP can successfully identify small groups (provided a small proportion can be identified using hashing based methods) due to its supervised classification approach.

A large number of cells (n=2582, >10% of the dataset) are assigned to the negative group by HTODemux, meaning that they can’t be assigned due to their hashing quality. It was previously identified that cells with low hashing counts (negative) also had low RNA quality, and so we next asked whether these ‘negative’ cells were truly low quality cells. Plotting library size and number of detected features for each negative cell, we see that the majority (2138 out of 2482, 86%) pass standard scRNAseq quality checks (Figure 4C). The SNP profiles of cells within and between demultiplexed groups can be visualised as a heatmap (Figure 4D). Visualising SNP profiles from the HTODemux Negative group, colouring cells by the HTODemux and demuxSNP classification, and splitting by the demuxSNP classification, a consistent SNP profile is observed in each reassigned group. Additionally, the majority of cells are reassigned to Hashtag5, consistent with the souporcell and demuxSNP annotations. Additionally, to validate the reassignments, we compare the Jaccard similarity between group-wise centroids of the reassigned “Negative” cells with those of cells from each hashtag group called with high confidence. Highest similarity is observed between reassigned cell groups and their corresponding high confidence singlet group with similarities of 0.97, 0.87 and 0.85 for Hashtag5, Hashtag3 and Hashtag4 respectively, the singlet groups to which the highest number of negatives cells were reassigned (Supplementary Figure 2A).

Examining binary distance distributions of singlet group SNP profiles, the presence of a multimodal or bimodal distribution may indicate cells from multiple biological samples (Figure 4E). Singlet groups where a bimodal distribution is evident tend to have fewer cells called the same by HTODemux and demuxSNP. Proportions of agreed cells between HTODemux and demuxSNP are 0.85, 0.70, 0.93 for Hashtags 1, 2, 4 compared to 0.94, 0.98 and 0.98 for Hashtags 3, 5, 6 respectively. Visualising the SNP profiles of HTODemux Hashtag2, the group with poorest agreement between HTODemux and demuxSNP, we can confirm multiple SNP profiles. The main SNP profile is called consistently as Hashtag2 by both HTODemux and demuxSNP. The remaining cells are reassigned by demuxSNP, mostly to Hashtag4 & Hashtag5 (Figure 4F). Again, we confirm the new annotations for this group against the high confidence annotations, noting specifically greatest similarity 0.99 and 0.95 for the main cell groups which were those that remained as Hashtag 2 (n=237) and those that were reassigned to Hashtag5 (n=74) respectively (Supplementary Figure 2B). Leveraging both SNP and hashing modalities, demuxSNP increases the number of assigned cells which would have otherwise been labelled as negatives, as well as reassigning cells misassigned due to hashing quality.

## Discussion

Multiplexing is primarily a cost reduction measure now utilised in most single-cell experiments, allowing full utilisation of high throughput assays. However, large numbers of negative, uncertain or misassigned cells resulting from suboptimal demultiplexing reduce its effectiveness. Accurate assignment of cells to their original source sample through demultiplexing is critical to interpretation of downstream analysis, minimising wasted data through misclassified or unclassified cells, as well as maintaining confidence in the technique as a cost saving measure to allow larger scale experiments. To this end, we make key contributions compared to existing hashing and SNPs based methods.

The dependence of hashing performance on hashing quality has been reported previously [6,15], yet current solutions to this problem have focused on more advanced modelling of the counts data to optimise detection of signal from noise. We proposed a novel method applicable to genetically distinct samples utilising cell hashing and SNPs, overcoming dependence on hashing data quality by assigning hashing negative or uncertain cells based on their SNP profiles. This results in overall improvements in classification performance, as well as assignment of hashing negatives and allowing the recovery of this valuable data. Application to real data shows that high RNA quality hashing negatives recoverable by demuxSNP may consist of over 10% of the total dataset. We are currently unable to estimate the expected distribution of hashing negatives across a large number of experiments and this measure for loss of data may be tracked in future benchmarking studies.

We additionally show systematic biases in genotype-free SNPs based method, souporcell. Despite being performant in many scenarios [20], its unsupervised classification results in systematic flaws when assigning cells to biological samples with few cells at high doublet rates, misclassifying doublets in place of the singlet group. While suggested as a potential limitation of a similar method previously [10], we believe this is the first time the problem has been described in greater detail. Our benchmarking results suggest a threshold of 40% doublets after which small groups are missed, however this may be dependent on other experimental factors and warrants future benchmarking. We go on to show that the demuxSNP supervised method shows increased robustness to doublet rate and class size imbalances. By subsetting the SNPs used, overall computational cost as well as data sparsity is reduced.

The majority of methods released to date leverage either SNP or hashing data. A recently described hybrid method HTOreader [21] uses results from a SNP based method such as souporcell to group cells that share a common genotype which are assigned to the dominant hashtag group in each genotype cluster. HTOreader assigns donor identity to cells that are assigned as singlet by either SNPs or cell hashing. The authors have demonstrated HTOreader’s usefulness for identifying samples missed by hashing methods. However, we expect any biases in the SNPs base methods, such as the behaviour described above where small groups are missed by genotype-free SNPs methods to be reflected in HTOreader’s results also (Supplementary Figure 3). This may be confirmed when full code becomes available.

We show that demuxSNP provides a framework for how both SNP and hashing data can be combined to optimise demultiplexing. As a result, an obvious limitation then exists that the method can only be applied to genetically distinct biological samples, and so cross validating demultiplexing results from genetically similar biological samples remains a challenge in the field. Other limitations of demuxSNP include dependence on the quality of the SNP data, and so reduction in sample sequencing depth or number of universally expressed genes result in more missing data and negatively impact performance. Similarly, the method performs poorly at classifying low RNA quality cells although this is less of a concern. While this applies to some extent to all SNP based demultiplexing methods, the KNN algorithm employed here is sensitive to missing data due to dropout and cell transcriptomic heterogeneity in scRNAseq. Future development using other supervised classification techniques that better handle missing and sparse data may extend this method.

More generally, benchmarking hashing demultiplexing methods in scRNAseq poses many challenges. Firstly, defining ground truth is non-trivial. In the absence of a tool to simulate realistic hashing data, SNP based methods have been used to define ground truth to benchmark hashing methods [6,7]. Consequently, any biases inherent in the SNPs based method will be reflected in the benchmarking results. Secondly, and following on from this, it is not feasible to generate sufficient benchmarking datasets to evaluate changes in experimental conditions such as doublet-rate, number of samples, sample imbalance This results in conclusions which are difficult or inappropriate to generalise, seen particularly in evaluating performance of the Seurat HTODemux function [6,7] due to limited benchmarking datasets. In contrast, for benchmarking of SNP based methods, strategies to simulate SNPs from real data have been developed elsewhere [14] and have been successful in evaluating the impact of factors such as doublet rate and ambient RNA content. This furthers the need for tools which allow realistic simulation of hashing counts, such as those used for simulating scRNAseq data [22] to make more comprehensive benchmarking studies feasible.

A recent development which acknowledges the role multiplexing plays in the future of single-cell studies was the release of the ‘multi’ functionality in Cell Ranger [23], the bioinformatics pipeline used to analyse data from the popular 10X Genomics single-cell platform using a Gaussian mixture model. While the exact model used has not, to our knowledge, been independently benchmarked, mixture model type methods have shown to be more consistently performant in this study and elsewhere. However, this update may come with some disadvantages. Previously, demultiplexing was carried out as part of downstream analysis, where the hashing quality could be reviewed, visualised and different algorithms tested. The incorporation of this step within the CellRanger pipeline will streamline downstream analysis but may consequently impede recovery of negative cells or identification of misassigned cells. Ongoing efforts to optimise laboratory protocols and workflows [24,25] to improve data quality or alternative labelling technologies [26] less susceptible to non-specific binding will be key in resolving this.

## Conclusion

Overall, we have shown that a multi-modal framework allows demuxSNP to recover hashing negative cells, reassign cells miscalled by hashing algorithms based on their SNP profile and overcome class size imbalance and doublet rate issues incurred by genotype-free SNPs methods while linking samples back to their group. The workflow has been implemented in the R/Bioconductor package [demuxSNP](https://bioconductor.org/packages/release/bioc/html/demuxSNP.html) providing additional functionality for assisting in SNP selection and selecting training data. The package provides interoperability with the Bioconductor SingleCellExperiment class.

## Methods

### demuxSNP workflow

1. SNPs are filtered to those located within genes expressed across most cells in the dataset
2. VarTrix uses filtered SNP list to call SNPs in each cell.
3. SNPs with few reads across cells in the dataset are removed.
4. Probabilistic hashing methods leveraged to determine high confidence singlets
5. Labels from high confidence singlets along with simulated doublets used to train KNN classifier and predict negative/uncertain cells.

To leverage classification techniques applicable to binary data, SNP status is recoded to absent/present (1,0) and k-nearest-neighbour classification (KNN) [27] is performed using Jaccard coefficient. The classification model parameters n (number of cells to downsample to) are estimated based on the training data size. To prevent bias in the training data due to class size, each class is downsampled to *n*, the smallest class size calculated from the training data, unless *n* < *nmin* in which case *nmin* is used for remaining groups above the threshold. Parameters *k* (number of nearest neighbours used in KNN classification) and *d* (coefficient of *n*, for which *d*n* doublets are simulated between each biological sample pairing default to values performant in most scenarios (Supplementary Figure 4). The optimum number of doublets simulated between singlet groups is approximately proportional to *n* and so can be described by *d*, a coefficient of *n* with optimal values ranging between 0.5-0.65. Similarly, sensible values of *k* are 25-50.

For the results shown, a starting list of SNPs from 1000 Genomes common variants occurring with frequency >5% [10] was used and filtered to those located within top 100 most commonly expressed genes. VarTrix [28] was used with default settings. Vartrix output was filtered to remove SNP locations not sequenced in the data (sequenced in <80% cells). High confidence cells were determined using demuxmix with acceptance threshold of 0.75. Classes of cells denoted as uncertain or negative were predicted using KNN. Parameter *n* was automatically calculated from the data. *d* and *k* were set at 0.5 and 25 respectively, as values shown to be robust over a large range of *n* and % doublets (Supplementary Figure 4.).

### Datasets

#### Single-cell RNA sequencing of ccRCC dataset

Single-cell RNA-seq experiments were performed by the Brigham and Women’s Hospital Center for Cellular Profiling. Sorted cells were stained with a distinct barcoded antibody (Cell-Hashing antibody, TotalSeq-C, Biolegend). After washing, the stained cells were resuspended in 0.4% BSA in PBS at a concentration of 2,000 cells per μL, then loaded onto a single lane (Chromium chip K, 10X Genomics) followed by encapsulation in a lipid droplet (Single Cell 5′kit V2, 10X Genomics) followed by cDNA and library generation according to the manufacturer’s protocol. 5′ mRNA library was sequenced to an average of 50,000 reads per cell, protein (hashtags) library sequenced to an average of 15,000 reads per cell, all using Illumina Novaseq.

#### Simulated datasets

##### Data preparation

Ground truth was obtained from multiplexed renal cell cancer experiment by applying demuxmix [17] with high acceptance threshold to generate a list of barcodes associated with each sample. From this, an individual bam file was generated per group using subset-bam [29] which forms the basis for SNP simulation.

##### SNP simulation

For the aligned reads, we followed the simulation strategy of Weber et al. [14] leveraging samtools [30]. Briefly, beginning with a single bam file per biological sample, a suffix is added to each cell barcode to identify cells from that group. The bam files are then merged. To simulate doublets, a lookup file is generated whereby randomly selected barcodes from a fixed number of cells are each reassigned to the barcode from a different cell.

##### Hashing simulation

Low quality hashing/uncertain cells are removed as part of the data preparation step. Using the same lookup file generated in the previous step, RNA and hashing counts for each doublet pair described in the lookup file are merged and the sum of their respective RNA and hashing counts are retained. To replicate low quality hashing data, the hashing signal is scaled down.

For the purposes of simulation, percentage doublets targeted and described in the manuscript include single-sample and multi-sample doublets. For the purposes of measuring demultiplexing performance, single-sample multiplets are considered as singlets and multi-sample multiplets considered doublets, as demultiplexing methods are only capable of identifying multi-sample doublets. Data simulation steps and analysis are incorporated into an adaptable and reproducible Nextflow [31] pipeline.

#### Benchmarking methods

souporcell [12] was applied to case study and simulated datasets using 1000 Genomes common variants generated by the authors of Vireo [10] and skipping remapping with default parameters.

cellhashR [15] ‘GenerateCellHashingcalls’ was used to accommodate use of multiple algorithms [4,15,17] and ensure consistency across preprocessing.

#### Computational Cost

‘date’ shell command was applied at start and end of process/workflow and the difference taken. Average compute time is taken over different % doublets and seeds. Each command was run on 10 cores of Intel(R) Xeon(R) Gold 6342 CPU

#### Renal cell cancer case study

souporcell was applied using default parameters and common variants supplied from 1000 Genomes common variants with >5% frequency.

Hashing data was normalised using the centred log ratio method from NormalizeData() and demultiplexed using HTODemux using default parameters.

Low quality cells were determined as those with fewer than 1,000 UMIs per cell and 500 genes per cell.

To evaluate demuxSNP’s reassignment in the absence of ground truth, the SNP profiles of reassigned cells are compared to those of cells called with high confidence by demuxmix. The Jaccard similarity between the centroid of each reassigned group and that of the corresponding high confidence singlet group is calculated. The reassigned cells compared are from the HTODemux negative (large number of unassigned cells) and Hashtag2 (suspected largest number of misassigned cells) group. Greater similarity across the leading diagonal is indicative of cells being correctly reassigned (Supplementary figure 2).

Visualisation of SNP profiles within and between groups provides a useful assessment of whether misassigned cells are present in the data. Plotted as heatmap, distinct SNP profiles may appear within groups. Quantifying the genetic variability within each assigned group also allows for assessment of the demultiplexing results. We used the vegdist function from the vegan [32] package to calculate the binary Jaccard distance between cells within the same assigned group. Homogeneous groups (containing mostly cells from a single sample) will appear unimodal whereas groups containing misassigned cells from different groups will be more heterogeneous and will appear as multimodal.

Plots were generated using ComplexHeatmap [33], ggpubr [34] and ggalluvial [35].

## Availability of Source Code and Requirements

Project name: demuxSNP

Project home page: https://doi.org/doi:10.18129/B9.bioc.demuxSNP

Operating system(s): Windows, MacOS, Linux

Programming language: R

Other requirements: VarTrix

License: GNU GPL

Project name: demuxSNP-paper-figures

Project home page: https://github.com/michaelplynch/demuxSNP-paper-figures

Operating system(s): Linux

Programming language: R, Shell

Other requirements: Slurm Workload Manager, Nextflow, Environment Modules, Apptainer, Conda

License: GNU GPL

## Data Availability

Raw and processed multiplexed sequencing data generated in this study (ccRCC dataset) are available from the Gene Expression Omnibus (GEO) (accession GSEXXXXXX).

## Abbreviations

scRNASeq: single-cell RNA sequencing
SNP: single nucleotide polymorphism
KNN: k-nearest neighbours

## Additional Files

Supplementary_figures.pdf

## Ethics Approval and Consent to Participate

Renal cell carcinoma specimens were collected under DFCI approved protocol #19-194 and #98-063.

## Authors’ contributions

M.P.L.: Conceptualization, Formal Analysis, Software, investigation, Data curation, Writing - Original Draft Preparation, Writing - Review & Editing, Visualisation

Y.W.: Resources, Investigation, Writing - Review & Editing

L.G.: Supervision, Writing - Review & Editing

A.C.C.: Conceptualization, Resources, Writing—Review & Editing, Supervision, Funding acquisition

## Acknowledgements

Prof. Wayne A. Marasco for use of data and seed networks team for discussions.

