## Supplementary Figures for "demuxSNP: supervised demultiplexing scRNAseq using cell hashing and SNPs"

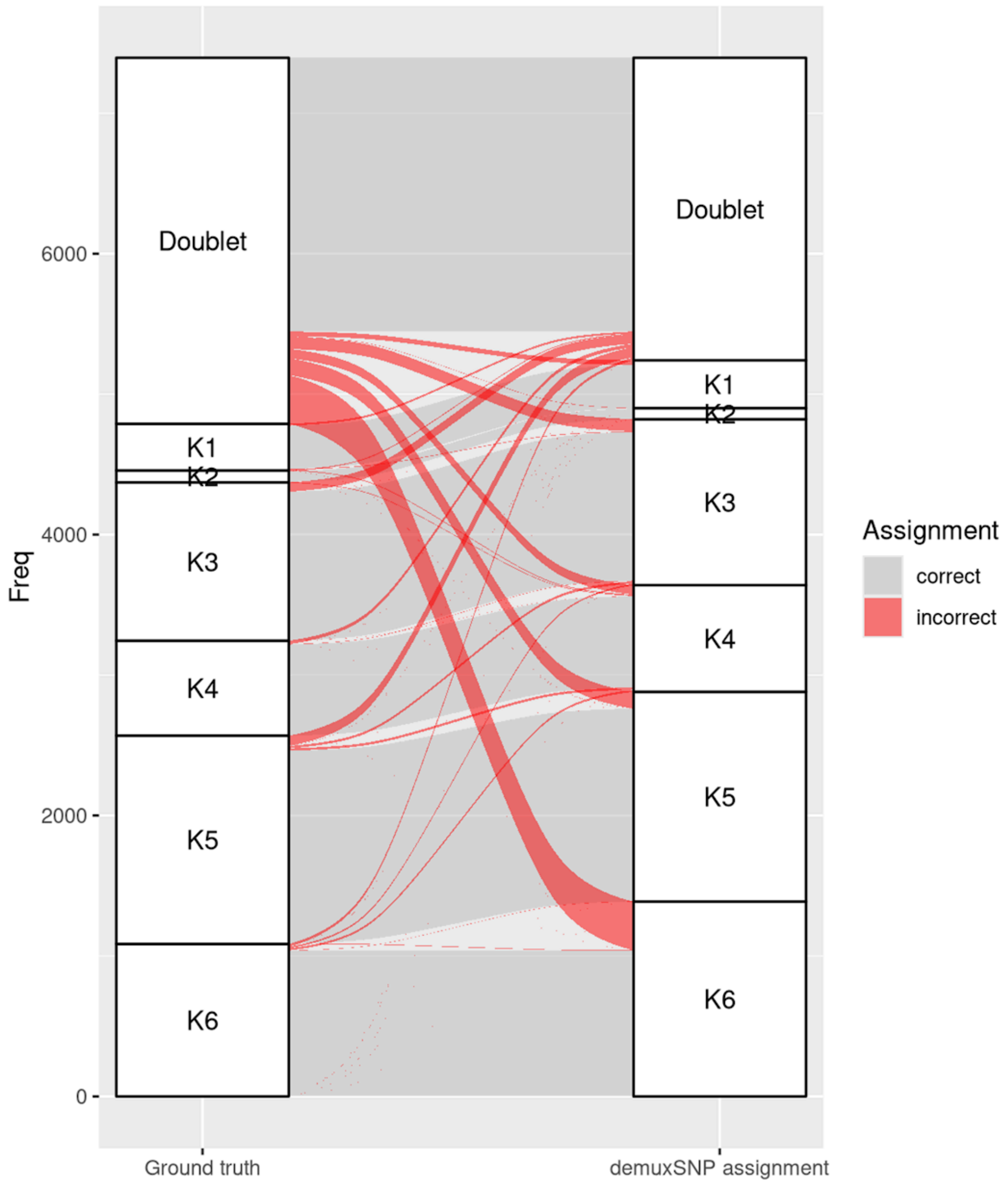


Supplementary Figure 1. demuxSNP correctly assigns K2 group.

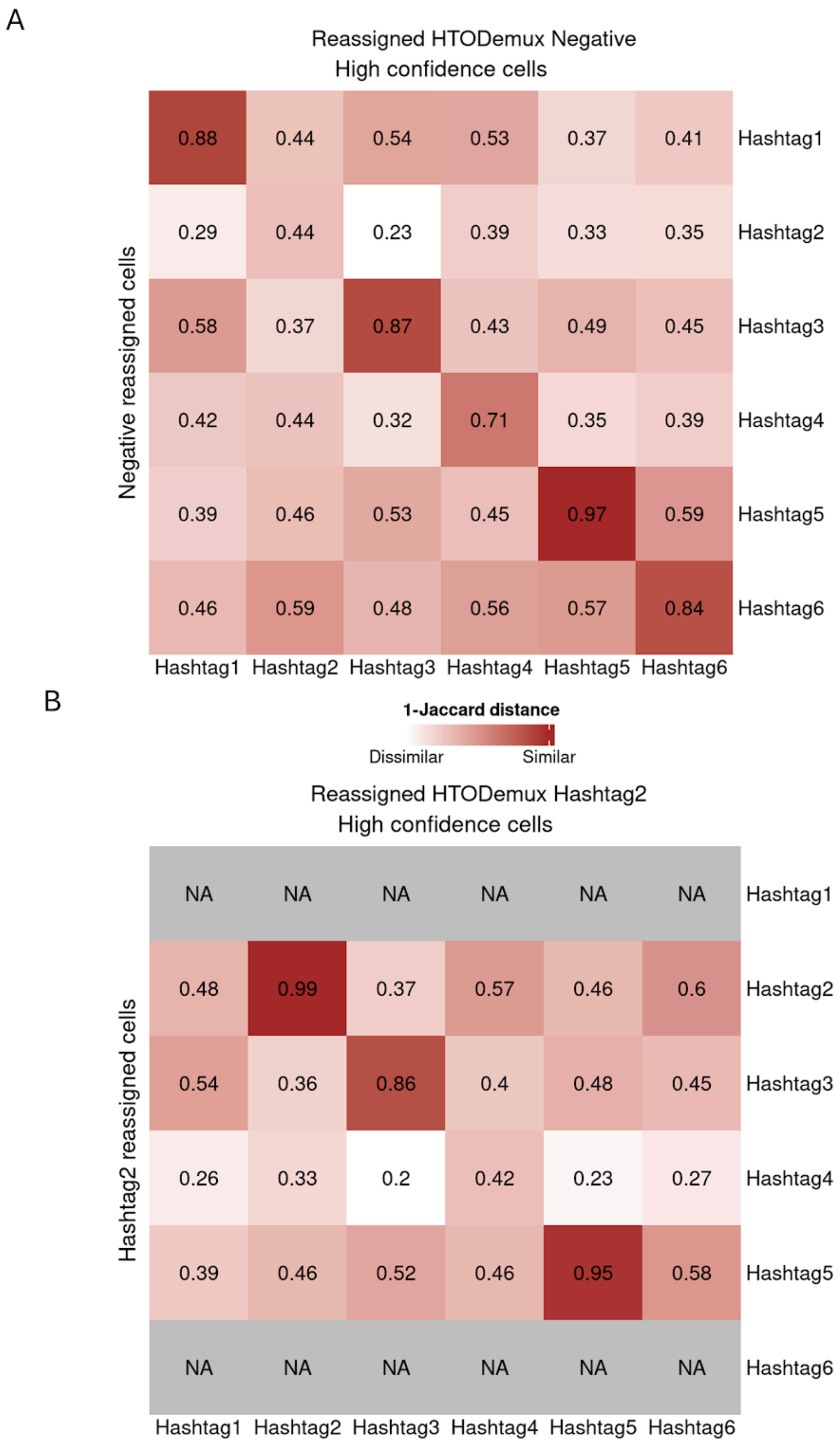


Supplementary Figure 2. (A) Similarity of HTODemux negative cells reassigned by demuxSNP to high confidence calls. (B) Similarity of HTODemux Hashtag2 cells reassigned by demuxSNP to high confidence cells.

Souporcell labels

| HTOreader labels | 2 | 3 | 4 | 5 | 1 | 6 | Doublet | unassigned |
|---|---|---|---|---|---|---|---|---|
| Hashtag1 | 1195 | 2 | 12 | 2 | 3 | 0 | 38 | 141 |
| Hashtag2 | 2 | 1 | 1 | 150 | 4 | 0 | 102 | 45 |
| Hashtag3 | 4 | 15 | 2755 | 0 | 5 | 0 | 42 | 228 |
| Hashtag4 | 3 | 7 | 11 | 1693 | 11 | 0 | 59 | 302 |
| Hashtag5 | 2 | 2 | 4 | 3 | 3152 | 0 | 33 | 125 |
| Hashtag6 | 2 | 6 | 1 | 3 | 2 | 0 | 43 | 91 |
| Doublet | 208 | 799 | 605 | 271 | 804 | 0 | 2817 | 195 |
| Negative | 58 | 16 | 120 | 30 | 150 | 0 | 26 | 59 |

Supplementary Figure 3. Confusion matrix between HTOreader and souporcell assignments.

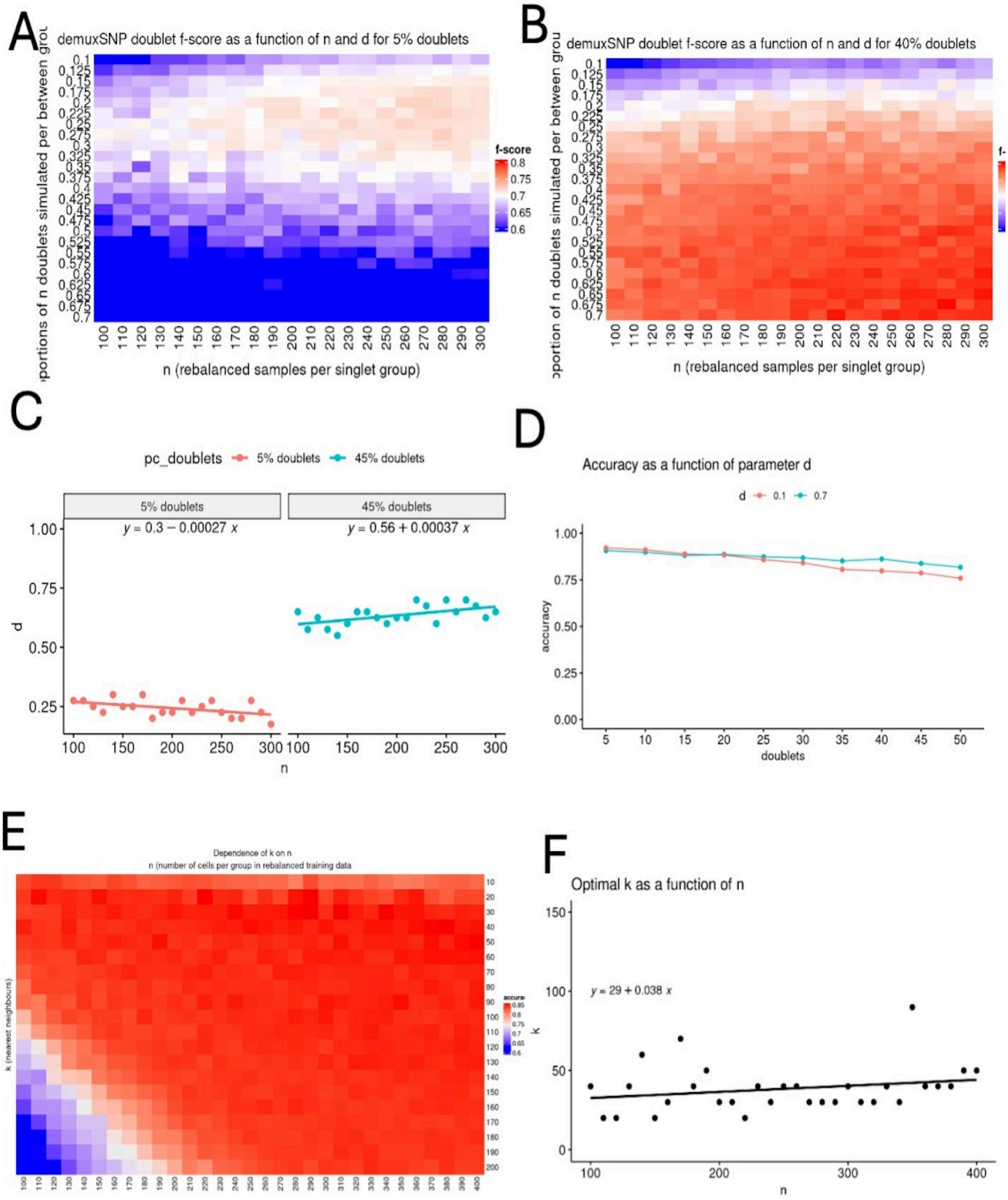


Supplementary Figure 4. Parameter choice for *k* and *d*. *(A) doublet f-score (average of five runs) varying d and n on low doublet rate data (5%). (B) doublet f-score (average of five runs) varying d and n on high doublet rate data (45%). (C) Linear model fitting optimal d to n. (D) Performance comparison on high d and low d. (E) Overall classification accuracy (average of five runs) decreases when k > n. (F) Linear model fitting optimal k to n suggesting suitable values between 25-50.*
